# Codominance of two symbiont genera within the same coral host is associated with elevated symbiont productivity and lower host susceptibility to thermal stress

**DOI:** 10.1101/2021.01.20.427463

**Authors:** Evelyn Abbott, Groves Dixon, Mikhail Matz

## Abstract

In recent years, as sea surface temperature increases, many coral species that used to harbor symbionts of the genus *Cladocopium* have become colonized with the thermally tolerant genus, *Durusdinium.* Here, we asked how the symbionts of one genus react to the presence of another symbiont genus within the same coral host, and what effect this interaction has on the host. We used previously published transcriptomic data from *Acropora hyacinthus* corals hosting *Cladocopium* and/or *Durusdinium* symbionts and looked at gene expression in all three symbiotic partners depending on the relative proportion of the two symbiont genera within the same host. We find that both *Cladocopium* and *Durusdinium* change their expression the most when their proportions within the host are nearly equal (the state that we call “codominance”): both genera elevate expression of photosynthesis and ribosomal genes, suggesting increase in photosynthesis and growth (i.e. higher productivity). At the same time, the coral host also elevates production of ribosomes suggesting faster cellular growth, and, when heated, shows less pronounced stress response. These results can be explained in two alternative ways. One explanation is that increased competition between codominant symbionts switches them to the higher productivity mode, which benefits the host, making it more resilient to stress. Alternatively, the symbionts’ elevated productivity might be not the cause but the consequence of the host being particularly healthy. Under this explanation, rapid growth of the healthy host creates new space, lowering the symbioints’ competition and thus promoting their growth, which allows for codominance to happen where one genus would otherwise outcompete another. The latter explanation is supported by the fact that codominance is associated with lower symbiont densities, assessed as relative proportions of symbiont reads in the data. Irrespective of the causation direction, the presence of mixed symbiont communities could potentially be used as an instant indicator of coral well-being, which would be a useful tool for coral conservation and restoration.

## Introduction

Reef building corals get up to 90% of their energetic requirements for growth and calcification through symbiosis with dinoflagellate algae of the family Symbiodiniaceae (Falkowski *et al.* 1984). Coral bleaching is the breakdown of this symbiosis, and often occurs when water temperatures exceed a certain threshold. This heat tolerance threshold varies more depending on the genus of symbiont than on the host genetics (Fuller *et al.*, 2020). These symbionts, formerly delineated as clades A-I, have now been divided into six genera (Lajeunesse *et al.*, 2018). In the Great Barrier Reef, the majority of scleractinian corals of the genus *Acropora* have historically engaged in symbiosis with symbionts of the genus *Cladocopium*. However, as ocean temperatures continue to rise and bleaching events become more frequent, these corals are becoming colonized by relatively opportunistic, thermotolerant symbionts of the genus *Durusdinium*.

Symbionts are known to have diverse effects on their coral hosts. Although *Durusdinium* symbionts confer bleaching resistance, it comes at the cost of reduced growth (Pettay *et a*l., 2015). Other physiological trade-offs have been observed, including reduced fecundity, reduced carbon acquisition (Matthews *et al.*, 2018), reduced calcification (Pettay *et al.*, 2015), and disease susceptibility (Shore-Maggio *et al.*, 2018). Furthermore, it was found that in the Caribbean coral *Montastraea cavernosa, Durusdinium* dominance is associated with differential expression of stress-related genes in the host: having *Durusdinium* appears to stress the host (Cunning *et al.*, 2020).

Most corals only associate with a single symbiont type at a time, with background levels of other symbionts present in host tissues (Baker, 2003). In the case of acroporid corals from the Great Barrier Reef, hosts may harbor *Cladocopium* symbionts with background levels of *Durusdiunium*, or the reverse, although some colonies have been shown to have a more even mixture (Ulstrup *et al.*, 2003). Whether these two symbiont genera interact in the host tissues is presently unknown. Furthermore, as colonization of acroporid corals by *Durusdinium* becomes more common, it is unknown how harboring two distinct symbiont genera at once impacts both the symbionts and the host.

In this study, we analysed existing gene expression data from two studies with a combined total of 181 *Acropora hyacinthus* samples (Rose *et al.*, 2015; Barshis *et al.*, 2013). These indo-pacific corals had entirely *Cladocopium,* entirely *Durusdinium*, or a mixture of both symbionts. We asked how symbionts respond to symbionts of a different genus within the same host. Initially, we predicted that symbionts would have the most distinct expression patterns depending on whether they are the majority or the minority within the host. We also anticipated that the stress of competition would cause the symbionts to become more virulent towards the host, prioritizing their own proliferation by sequestering more nutrients and translocating fewer photosynthates to the host (Lesser *et al.*, 2013; Baker *et al.*, 2018; Morris **et al.**, 2019). Therefore, we expected that corals hosting mixed symbiont populations would be more susceptible to heat stress and might show elevated expression of generalized stress response genes (Dixon *et al.*, 2020) even under non-stressful temperature. We were surprised to find no support for any of these predictions.

## Materials and Methods

### Data sources

We chose two studies (Rose *et al.*, 2015; Barshis *et al.*, 2013) for this analysis by searching for the genus, *Acropora,* in the NCBI SRA database, comprising 181 coral samples sequenced with RNA-seq method. These studies were selected because they involved similar heat stress experiments on *Acropora hyacinthus* corals. Furthermore, these corals were all isolated from back reef tide pools in Ofu, American Samoa. The SRA metadata tables for each study are shown in supplemental table S1.

### Sequence data processing and symbiont genus determination

Detailed descriptions of the data processing pipeline are on Github (https://github.com/evelynabbott/codominant_symbiosis.git). The Fastq files from both studies were downloaded using the SRA toolkit. Adapter trimming was done on paired-end mode using cutadapt, with a minimum length of 20 bp and a PHRED quality cutoff set to 20. FASTQC (Andrews, 2010) was used to assess the quality of a subset of 10,000 reads before and after trimming. Reads were then mapped to a combined reference comprising *Cladocopium* transcriptome*, Durusdinium* transcriptome (Ladner *et al.*, 2012), and *Acropora millepora* genome (Fuller *et al.*, 2018) using bowtie2. The reads in the resulting sam files were then split into three separate sam files, one for each organism. PCR duplicates were removed after alignment using MarkDuplicates from the Picard Toolkit (Broad Institute, 2019). Samtools (Li et al., 2009) was used to sort and convert from sam files to bam files. FeatureCounts (Liao, Smyth, & Shi, 2014) was used to count reads mapping to annotated gene boundaries.

### Differential gene expression analysis

We used DESeq2 to identify genes which were differentially expressed due to symbiont codominance and heat treatment. This analysis was performed on *Cladocopium* symbionts, *Durusdinium* symbionts, and the coral host. Genes with a mean count less than three were excluded from this analysis, leaving a total of 19,580 genes for the coral host, 15,332 for *Cladocopium*, and exactly 15,000 for *Durusdinium*.

### Weighted gene coexpression network analysis

This analysis (WGCNA) (Langfelder & Hovarth, 2008) was used to identify groups of co-regulated genes in both symbiont genera, to explore major patterns of gene expression variation in an unsupervised way. The input for this analysis were the matrices of normalized variance stabilized counts obtained with R function DESeq2::vst(), from which the variation due to study and to the logarithm of total read count was removed using the R function limma::removeBatchEfect. For *Cladocopium*, we ran WGCNA with a soft threshold power of 10, a minimum module size of 30, and a module merging threshold of 0.4. For *Durusdinium*, we ran WGCNA with a soft threshold power of 12, a minimum module size of 30, and a module merging threshold of 0.4.

### Functional enrichment tests

We used a Gene Ontology (GO) enrichment analysis that utilizes the Mann-Whitney *U* (MWU) test (Wright et al., 2017) to identify significant functional differences among up and down-regulated genes associated with the codominant state. We used −log10 transformed *p*-values output from DESeq2, multiplied by −1 when the gene was down-regulated, as input for this analysis, following (Dixon **et al.**, 2015). This test compares transformed p-value ranks among genes to determine whether the ranks of genes in a GO category diverge significantly from ranks of other genes. We also used a Eukaryotic Orthologous Group (KOG) MWU test (package KOGMWU in R, Dixon et al., 2015) to broadly determine coral host gene expression response to codominant symbionts. For this analysis, the data from this study was compared to the generalized stress response (GSR) profile from (Dixon et al., 2020) that represents the response of *Acropora* sp. to any kind of high-intensity stress.

## Results

### WGCNA

We were firstly interested in which factors impacted the gene expression of *Cladocopium* and *Durusdinium* symbionts the most. To find this out we identified groups of genes (“modules”) that were co-regulated across samples using weighted gene co-expression network analysis (WGCNA). These modules, which WGCNA identifies in an unsupervised fashion without knowledge of the experimental design, are then examined for correlation with known traits or experimental treatments. In this way, it is possible to identify the largest and most responsive groups of genes and then investigate what biological effects they track. By examining the modules’ behavior across samples (Fig. 1) we observed that the greatest gene expression difference in the symbionts was not between “genus-background” and “genus-dominant” states, as we initially expected. Instead, in both genera the most distinct state was when the proportions of the two symbionts were near equal within the host (Fig. 1 b,c), the state that we here term “codominance.”

**Figure 1.**
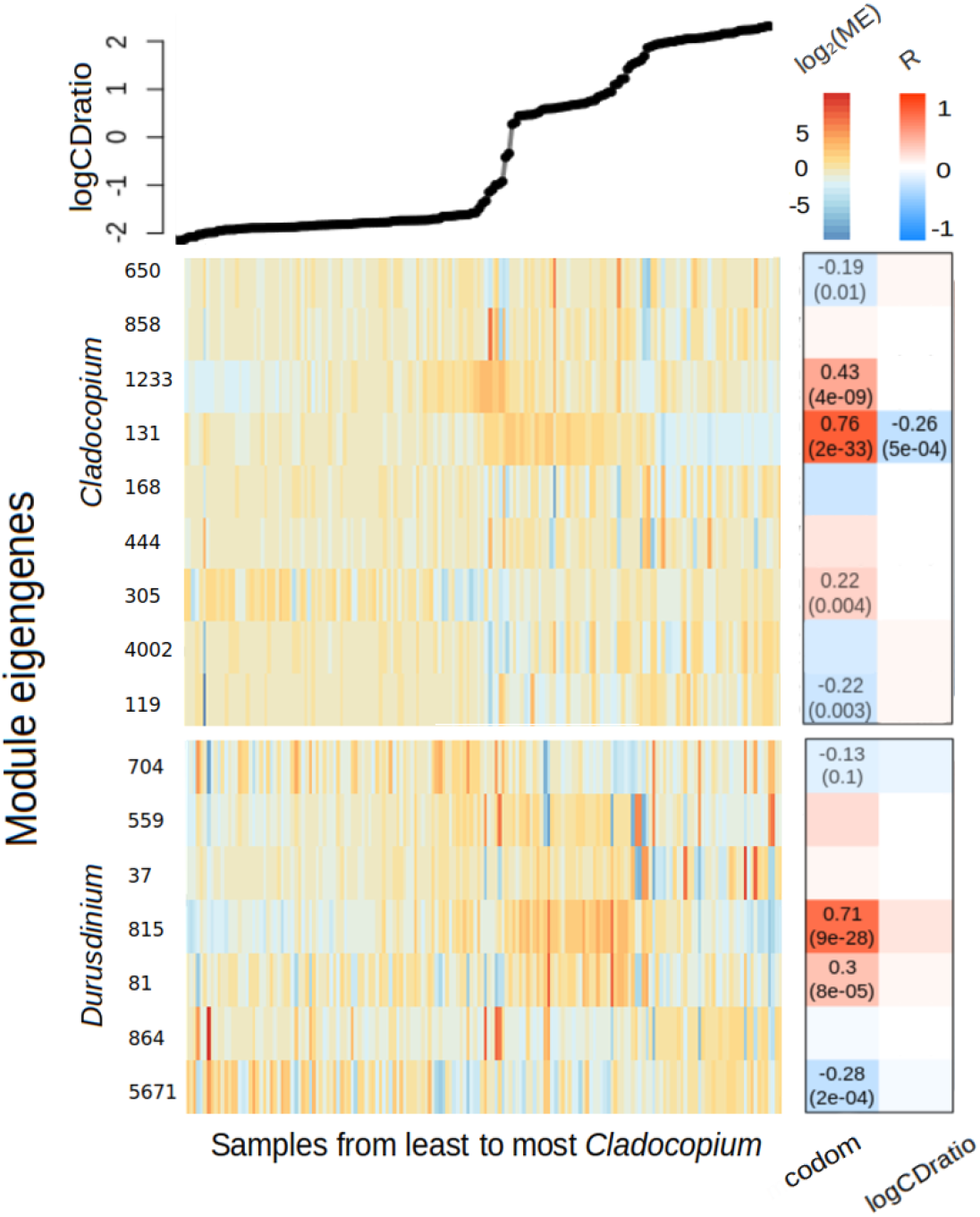
Correlation of WGCNA modules with symbiont codominance. Samples are represented across the x axis, in order from least to most *Cladocopium.* The points in the top panel show the log ratio of C:D counts for each sample. The heatmap rows represent WGCNA modules, with *Cladocopium* modules on the top and *Durusdinium* modules on the bottom; color representing the module’s eigengene expression (the left color scale). The numbers listed to the left of the heatmaps represent the number of genes in the module. The heatmaps on the right show Pearson correlations for each module with codominance (how near is the C:D ratio to 50:50, “codom”) and with the C:D ratio itself. Top number is the correlation coefficient and bottom number is p-value (only shown for significant correlations).

For *Cladocopium,* WGCNA identified nine modules of co-regulated genes. Two of these modules, containing 1233 and 131 genes respectively, were significantly correlated with codominance (r = 0.43, p < 4e-09 & r = 0.76, p < 2e-33). Additionally, the module containing 131 genes was negatively correlated with the dominance of *Cladocopium* (r = −0.26, p < 5e-04). For *Durusdinium*, WGCNA identified 7 modules of co-regulated genes. Two of these modules, containing 815 and 81 genes respectively, were significantly correlated with codominance (r = 0.71, p < 9e-25 & r = 0.3, p < 8e-05). One module, containing 5671 genes was negatively correlated with codominance (r = −0.28, p < 2e-04). Notably, neither symbiont had modules significantly associated with exposure to elevated temperature.

### DESeq2 analysis

We used DESeq2 modeling to measure the gene expression responses to codominance in each genus. Codominance was coded as a quantitative trait: log_10_(C:D ratio) multiplied by −1 when it was >0. This measure reaches its maximum (zero) when symbionts are equally represented and diminishes when their relative proportions become unequal. For *Cladocopium,* out of a total of 15,332 genes assayed, 1,930 were upregulated (false discovery rate, FDR = 0.1) and 89 were downregulated (FDR = 0.1, Fig. S1a). For *Durusdinium*, out of a total of 15,000 genes, 2,629 were upregulated (FDR = 0.1) and 2,868 were downregulated (FDR = 0.1, Fig. S1b). For the coral host, out of a total of 19,580 genes, 138 were upregulated (FDR = 0.1) and 83 were downregulated (FDR = 0.1, Fig. S1c).

We also evaluated gene expression response to heat treatment, and codominance:heat interaction (i.e., additional response to heat treatment under codominance), for all three partners. Response to heat was virtually absent in both symbiont genera: for *Cladocopium,* out of a total of 15,332 genes, 0 were upregulated and 1 was downregulated (FDR = 0.1, Fig. S2a), and for *Durusdinium*, out of a total of 15,000 genes, 1 was upregulated (FDR = 0.1) and 2 were downregulated (FDR = 0.1, Fig. S2b). The coral host, however, did show a pronounced response to heat treatment: out of a total of 19,580 genes, 1,399 were upregulated (FDR = 0.1) and 610 were downregulated (FDR = 0.1, Fig. S2c).

### Gene Ontology Analysis

To ascertain the functions of differentially expressed genes of symbionts in the codominant state, we used a Gene Ontology (GO) enrichment analysis that utilizes the Mann-Whitney *U* test (Wright et al., 2017). For *Cladocopium* symbionts, there were 2,019 differentially expressed genes (DEGs), with an FDR cut-off of 10%. At the GO level, the most notable signal pertained to ribosomes and photosynthesis. There were nine cellular component GO terms that were significantly (adjusted p < 0.001) enriched by upregulated genes. Of these, six were related to ribosome, and three to chloroplast (Fig. 2a).

**Figure 2.**
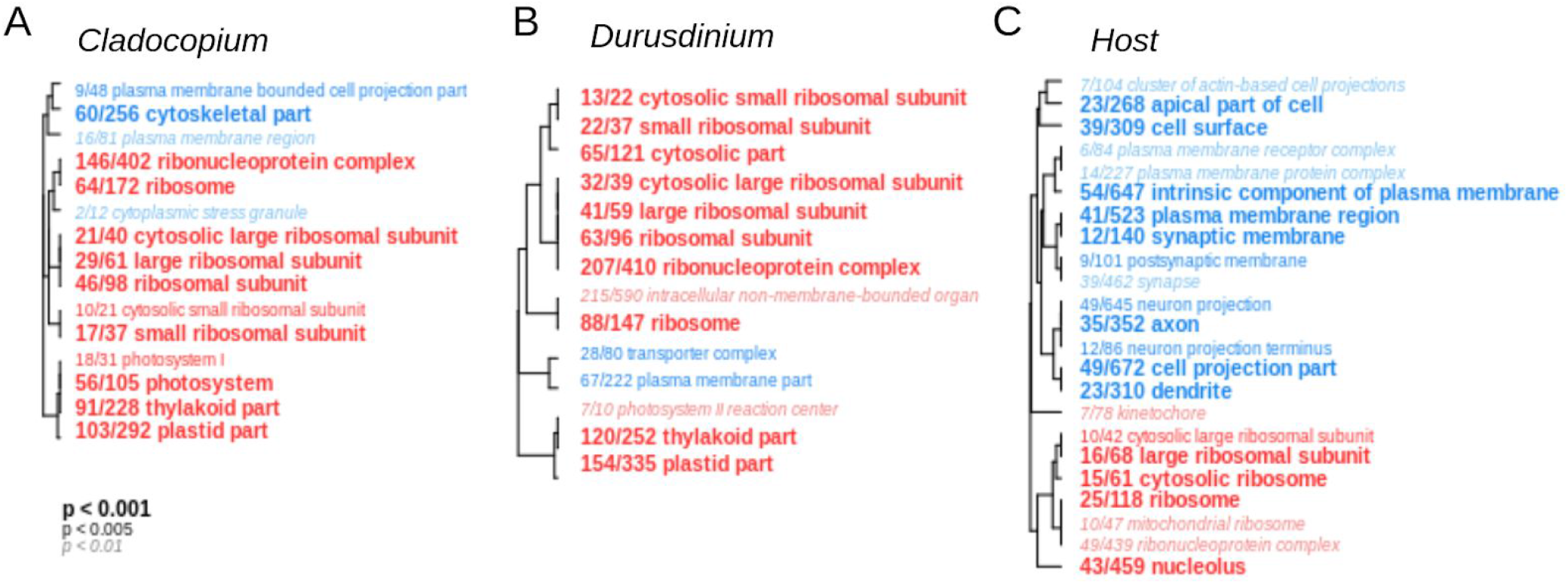
Gene ontology terms (cellular component) significantly enriched with up-regulated (red) or down-regulated (blue) genes based on Mann-Whitney *U* test for (A) *Cladocopium*, (B) *Durusdinium*, and (C) the coral host. Font indicates the *p*-value) adjusted for multiple testing over GO categories (see legend under panel A). Dendrograms represent hierarchical clustering of GO categories based on sharing of the genes among them. The fraction shows the number of genes with unadjusted *p*<0.05 in the DESeq2 model (numerator) relative to the total number of genes annotated with a given term (denominator).

We repeated the same analysis for *Durusdinium* symbionts, identifying 5,497 DEGs at the 10% FDR level. Like *Cladocopium*, the *Durusdinium* GO analysis results in multiple terms related to ribosomes and photosynthesis. There were ten cellular component terms that were significantly (adjusted p < 0.001) enriched by upregulated genes. Of these, seven were related to ribosomes and two to chloroplasts (Fig. 2b).

Lastly, we did this analysis for the coral host, identifying 7,799 DEGs at the 10% FDR level. Like the symbionts, there were multiple GO terms related to ribosomes. There were four cellular component terms that were significantly (adjusted p < 0.001) enriched by upregulated genes. Of these, three were related to ribosomes (Fig. 2c). Additionally, there were 11 GO terms significantly (adjusted p < 0.005) enriched for downregulated genes. Of these, six were related to neurons.

### Comparison of GO term delta-ranks between symbiont genera

To compare the functional signals of *Cladocopium* and *Durusdinium* symbionts in response to codominance, we used the GO_MWU analysis results which output the degree of up- or down-regulation of GO terms as measured by the “delta-rank” – the difference between the median rank of the genes annotated with the term and the median rank of all other genes. In GO_MWU, a positive delta-rank implies up-regulation of genes annotated with the GO term. The comparison of delta ranks (rather than per-gene expression changes) between experiments integrates the signal over meaningful functional categories of genes and, importantly for the current study, circumvents the problem of different reference transcriptomes used. At the GO delta-rank level, responses of *Cladocopium* and *Durusdinium* to codominance were clearly similar (Fig. 3), although the statistical significance of this similarity cannot be formally ascertained because GO terms are not independent (they may encompass overlapping sets of genes). Most notably, ribosomal and photosynthesis categories were similarly and significantly (p < 0.001) enriched for upregulated genes in both genera. These GO categories include large ribosomal subunit, small ribosomal subunit, ribonucleoprotein complex, thylakoid part, photosystem, and plastid part (p < 0.001).

**Figure 3.**
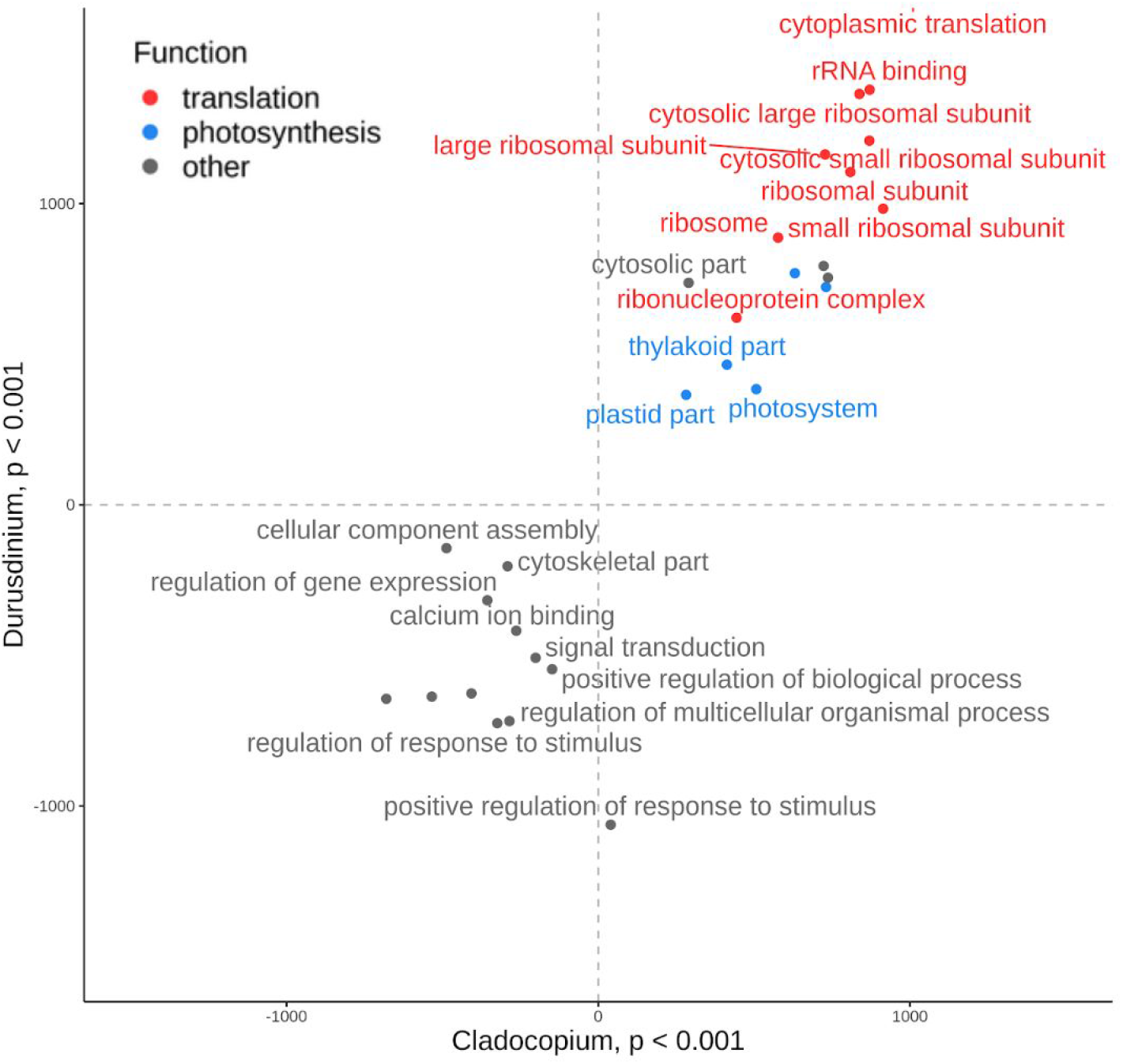
Delta rank comparison of highly significant (p < 0.001) *Cladocopium* and *Durusdinium* GO categories. Higher delta ranks indicate upregulation and lower delta ranks indicate downregulation.

We were concerned that the variance in the number of symbiont counts per sample would skew the results of our functional analyses. This was primarily due to the fact that read counts for each symbiont covaried with the symbiont’s proportion in the host. While this effect was formally accounted for in our analyses by including the log10(total count) as a covariate in DESeq2 models, doubts remained as to whether this would not create additional artifacts. One reassuring argument is that the main gene expression response (to codominance) was not aligned with the minor-major axis (i.e. with variation in total counts) but rather happened in the middle of the total counts’ scale. Still, to further test for possible residual effects of total counts variation on our results, we resampled the counts across samples so that each sample would have similar counts. We reduced the counts to 30,000 per sample, 15,000 per sample, and 8,000 per sample, the latter being the lowest number of counts for a symbiont type observed in a sample. We repeated the DESeq2 and GO_MWU analyses for each resampled dataset and compared the GO delta ranks of each. Reassuringly, we observed similar results at the GO delta-ranks level across all resampling trials, indicating that our functional results were largely unaffected by differences in coverage across samples (Fig. S3).

### Symbiont codominance is associated with reduced coral stress

We were next interested in how symbiont codominance is reflected in the host functional profile. In addition to analyzing the effect of codominance on its own, we were also interested in how symbionts’ codominance modifies the host response to heat stress (i.e., codominance:heat interaction). To better interpret the coral host functional profiles, we used eukaryotic orthologous group (KOG) analysis to identify broader functional categories of modulated genes and to formally compare these results to the known profile of *Acropora* sp. response to various kinds of high-intensity stress (generalized stress response, “GSR”; Dixon *et al.*, 2020). This analysis revealed that host gene expression response to symbiont codominance had no significant correlation with the GSR (Fig 4B), indicating that harboring codominant symbionts does not stress the host. Instead, supporting the GO analysis results, the “translation, ribosomal structure, and biogenesis” category was up-regulated (Fig. 4A), suggesting stimulation of cellular growth (Giordiano **et al.**, 2015; Elser *et al.*, 2003; Bosdriez **et al.**, 2015; López-Maury *et al.*, 2008). As expected, gene expression of the heated corals strongly correlated with the GSR (r = 0.66, p = 0.00058, Fig. 4C), implying that the corals were stressed by the treatment. However, delta-ranks for the heat:codominance interaction term were negatively correlated with the GSR (r=−0.79, p=5.9e-06, fig. 4D), which means that hosting codominant symbionts mitigated the heat stress response.

**Figure 4.**
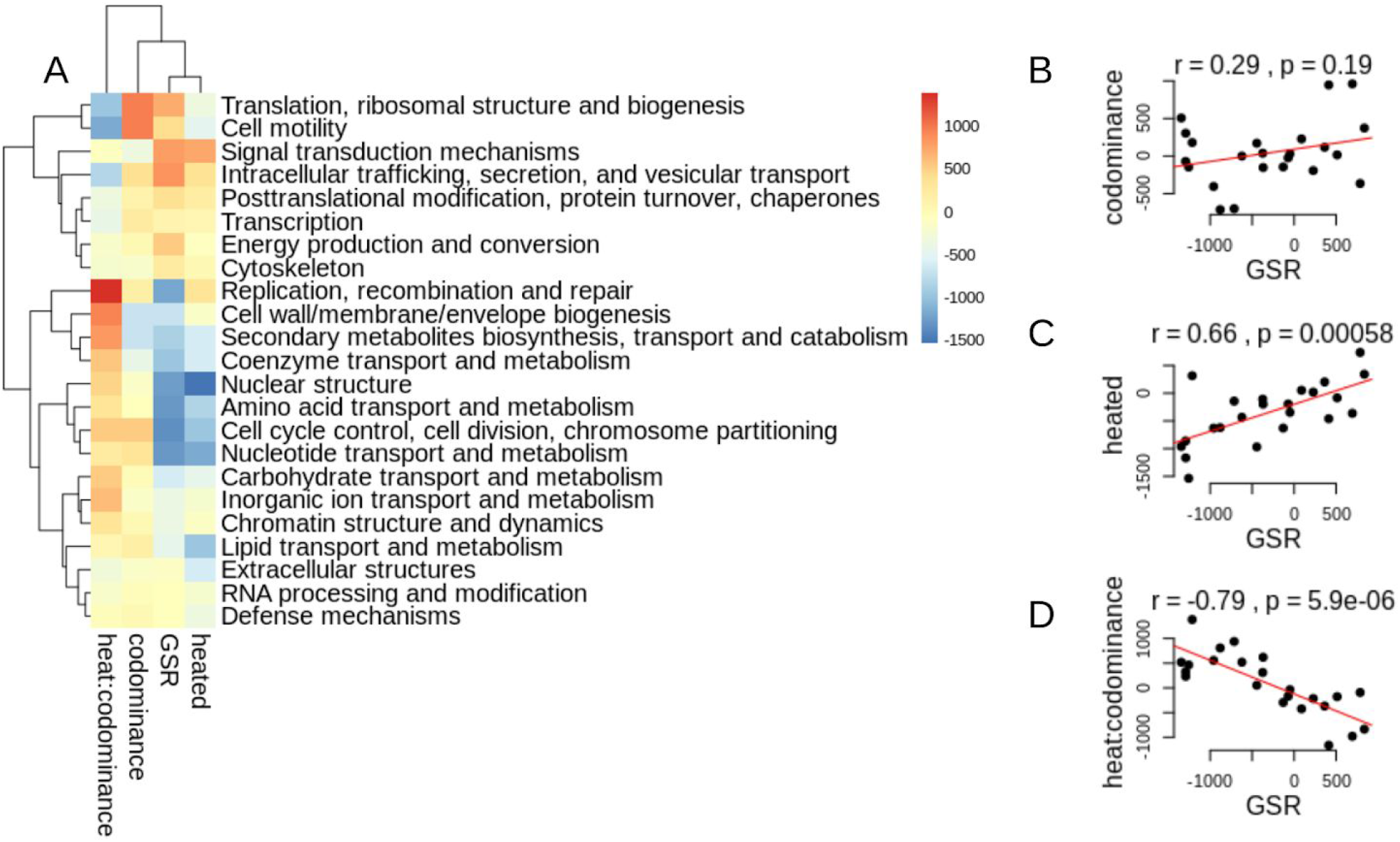
Symbiont codominance does not cause host stress and is associated with less pronounced host stress under heat. (A) Heatmap of delta-ranks across eukaryotic orthologous groups (KOGa), showing which KOG terms are upregulated (red) or downregulated (blue) with respect to heat:codominance interaction, codominance, generalized *Acropora* stress response (GSR), and response to heat in this experiment (“heated”). (B-D) Correlation of KOG delta ranks between generalized stress response (x-axis) and (B) response to codominance; (C) response to heat; (D) heat:codominance interaction (i.e., codominance-specific modification of the response to heat).

### Symbiont densities are the lowest under codominance

To evaluate the relative density of symbionts across samples, we looked at the variation of the ratio of symbiont to host reads. In another coral, *Orbicella faveolata*, this measure is a reliable proxy of symbiont density (Manzello *et al.*, 2018). We find that the proportion of symbiont reads is the lowest under codominance (Fig. 5).

**Figure 5.**
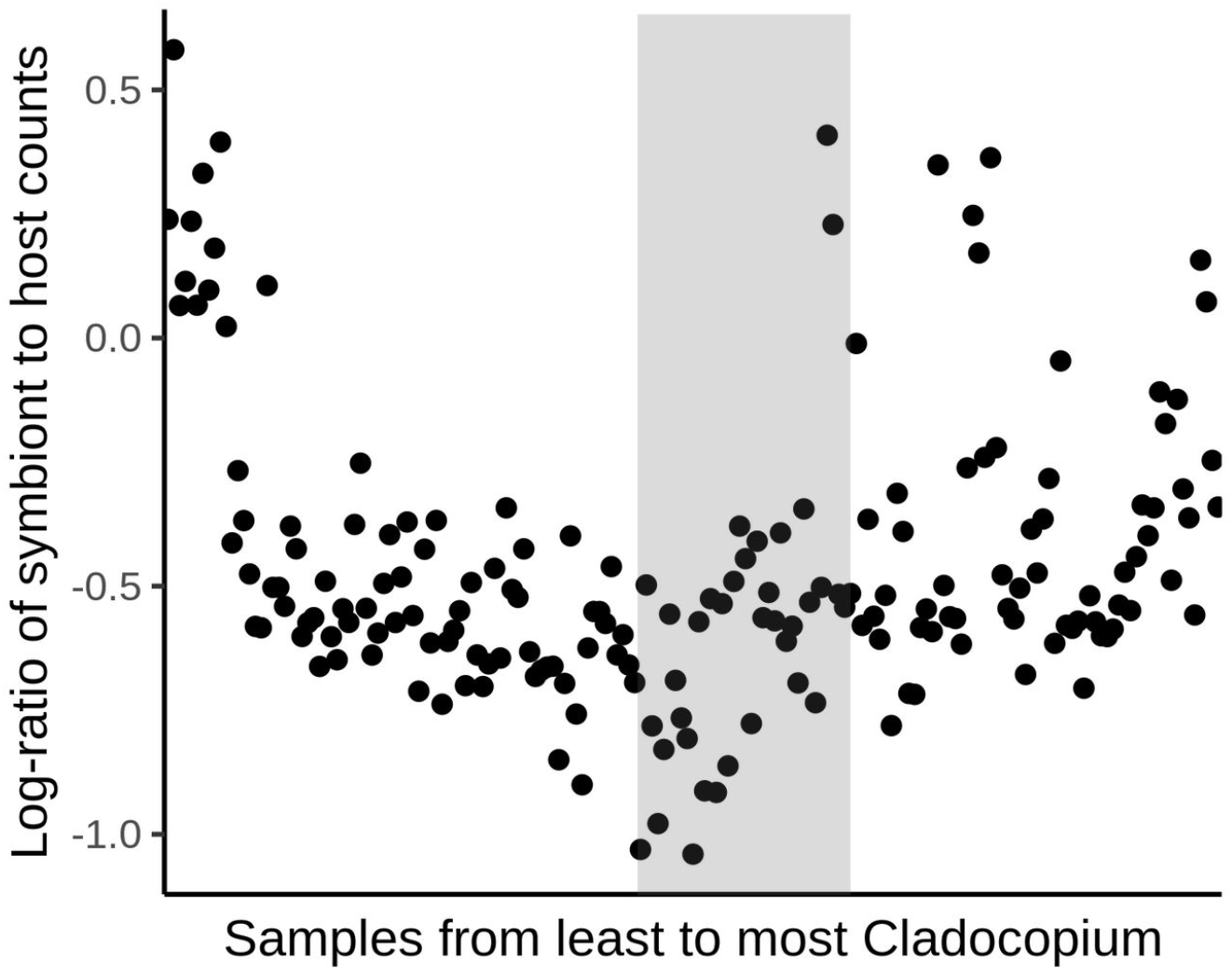
Corals with codominant symbionts tend to have lower symbiont density. X-axis is the rank of the sample from least to most Cladocopium, as on the top panel of Figure 1. The codominance zone is shaded. Y-axis is the logarithm of the ratio of the total number of reads mapping to symbiont transcriptomes and the number of reads mapping to the chromosome 1 of the host.

## Discussion

### Symbionts

We initially expected that symbiont gene expression would primarily depend on its own abundance within the host. Instead, in both *Cladocopium* and *Durusdinium*, gene expression was similar when the genus was either the overwhelming majority or minority, but was highly distinct when the relative proportion of the two genera were close to equal (codominant state, Fig 1). Functional analysis of the codominant state revealed upregulation of translation and photosynthesis machinery in both symbiont genera (Fig. 2, 3). Association between high growth rate and high concentrations of ribosomes has been demonstrated in a variety of organisms, including multicellular plants, green algae (Giordiano *et al.*, 2015), insects, crustaceans (Elser *et al.*, 2003), bacteria (Bosdriez *et al.*, 2015), and yeast (López-Maury *et al.*, 2008). We therefore believe that the observed functional signal indicates higher growth rate in both symbiont genera in the codominant state.

### Coral host

Under heat stress, symbionts are expected to parasitize the coral host by sequestering host resources and proliferating in host tissues without giving photosynthates to the host (Lesser *et al.*, 2013; Baker *et al.*, 2018; Morris *et al.*, 2019). Higher competition between symbionts typically results in higher virulence towards the host (Bremermann and Pickering, 1983; Chao *et al.*, 2000). In the codominant state the symbionts might be expected to compete more, and, as our data indicate, they also grow more, potentially withholding resources from the host. All this could result in host stress, but our data do not support this prediction. Functional profiles of corals with codominant symbionts did not significantly correlate with profiles from stressed corals (Fig. 4 A, B). Instead, under codominance the corals upregulated their translational machinery (Fig. 3 C, Fig. 4 A), which might be an indication of higher growth rate (Elser *et al.*, 2003; López-Maury *et al.*, 2008; Giordiano *et al.*, 2015; Bosdriez *et al.*, 2015). Down-regulation of neuronal components (Fig. 4 A) is also notable, but cannot be easily interpreted. More importantly, in corals with codominant symbionts the response to elevated temperature treatment was reduced (Fig. 4 D), indicating higher stress resilience.

### Causation direction

Two alternative hypotheses can be put forward to explain these results, differing in the direction of the causation. The first explanation is that symbionts switch into a highly competitive physiological mode when they are near 50:50 ratio in the host, increasing their productivity in the effort to outgrow each other. This “growth race” benefits the host because of higher symbiont productivity, improving the host’s growth and boosting its stress resilience. The second explanation starts with a host that is very healthy: it is less susceptible to stress and experiences a high growth rate. The host’s growth promotes symbiont growth to occupy the newly available space, while reduced competition inside the new space allows for symbiont codominance where one genus would otherwise outcompete the other. In support of this latter hypothesis, corals with codominant symbionts had lower symbiont densities than non-codominant corals (Fig. 6); this result would have been the opposite if the codominance was associated with the higher competition between symbionts, as postulated under the first explanation. The second explanation also appears more parsimonious because it does not assume an unknown mechanism of the two symbionts sensing each other’s abundances. In addition, the notable reduction in neuronal investment in codominant corals (Fig. 2C) suggests that codominance might in part be promoted by reduced host control over the symbionts’ proliferation, assuming host neurons are actually involved in such control, which remains to be investigated in the future.

## Conclusions

We have documented a strong gene expression response to the presence of a mixed symbiont community in both the symbionts and the coral host. Overall, the presence of a mixed symbiont community is associated with higher physiological fitness of all symbiotic partners involved, manifested as higher growth rate and productivity in symbionts and higher cellular growth and stress resilience in the host. It appears more likely that symbiont growth and productivity are elevated as a consequence of higher host fitness, not the other way around, which is a testable hypothesis for future research. Irrespective of the causation direction, the presence of mixed symbiont communities could potentially be used as an instant indicator of coral well-being, which would be a useful tool for coral conservation and restoration.

## Supplemental

**Supplemental Fig S1.**
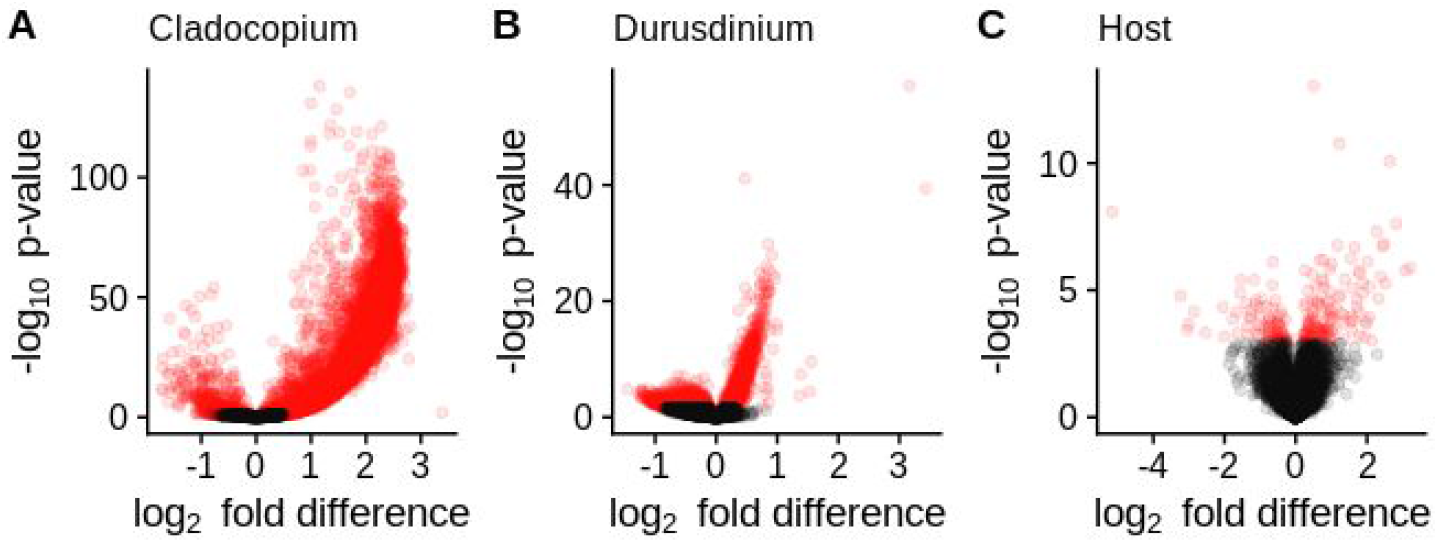
Volcano plots of codominance-associated gene expression changes for (A) *Cladocopium*, (B) *Durusdinium,* and (C) the coral host. Each point represents a gene. Red points represent FDR < 0.1.

**Supplemental Fig S2.**
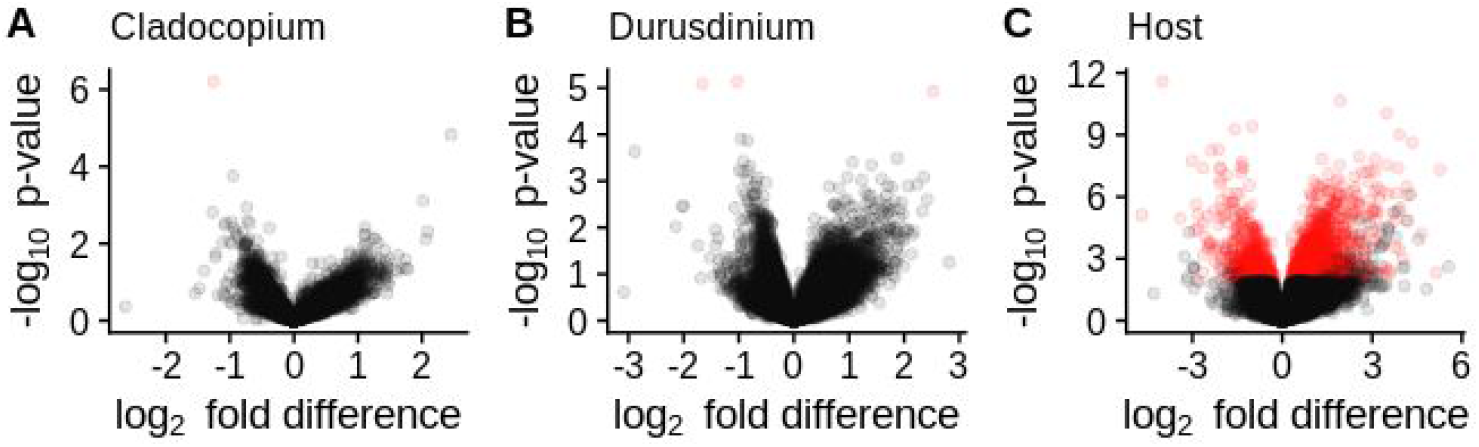
Volcano plots of gene expression changes in response to heat, for (A) *Cladocopium*, (B) *Durusdinium,* and (C) the coral host. Each point represents a gene. Red points represent FDR < 0.1.

**Supplemental Fig S3.**
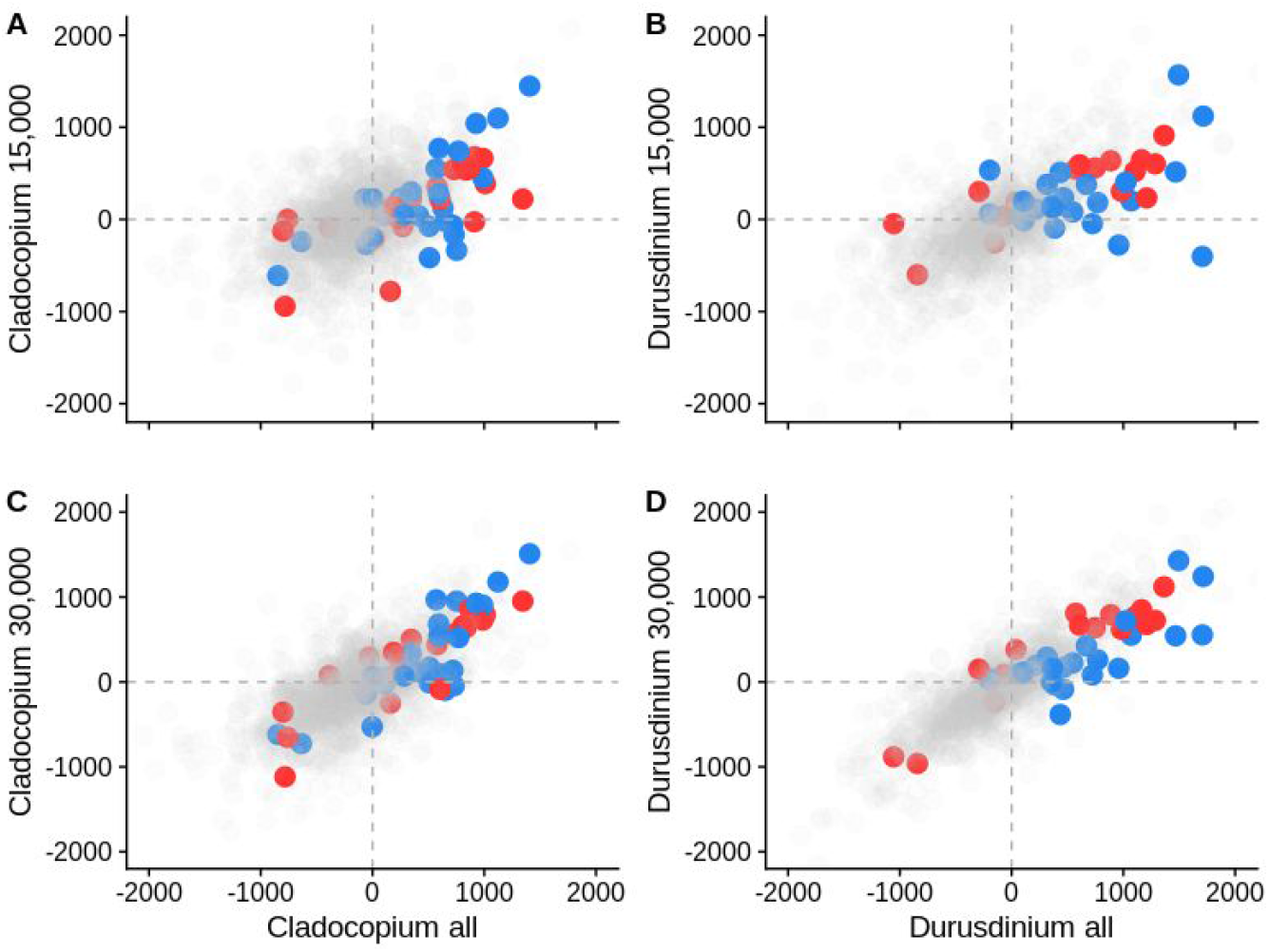
Functional signal in response to codominance is robust to low gene counts. GO delta-ranks for the full dataset (X-axis) are compared to GO delta-ranks obtained with subsampled data (Y-axis). A, B - subsampling to 15,000 counts per sample; C, D - subsampling to 30,000 counts per sample. A, C - *Cladocopium*, B, D - *Durusdinium*.

